# Evaluation of the antimicrobial activity of the alcoholic extract of *Tagetes erecta* against bacteria and fungi of clinical importance

**DOI:** 10.1101/2024.09.27.615189

**Authors:** Gabriel Martínez González, Frida Suárez González, Alejandro Chávez Alanuza, Sandra Stephanie Montes de Oca Cuenca, Angélica Guadalupe Cuarto Hernández, Lidia de Jesús Olvera, Jorge Angel Almeida Villegas

## Abstract

Antibiotic resistance is a significant global health concern, putting at risk treatment options for infectious diseases. As a result, treatments can be expensive and ineffective, so it is essential to look for new options such as the use of plant extracts with antimicrobial properties to combat these diseases. *Tagetes* sp is a plant species that is used in folk medicine to cure various diseases and has several properties among which the antimicrobial, for this reason, the aim of this research was to evaluate the antimicrobial activity of the ethanolic extract of *Tagetes erecta* L against various bacteria and yeasts. The antimicrobial activity of the extract was analyzed by the Kirby-Bauer method at different concentrations. The extract showed antimicrobial activity by presenting inhibition halos mainly against *Staphylococcus*, obtaining the following results: *S. aureus* ATCC 43300 (19,400 ± 0.435 mm to 500 mg/mL); *S. aureus* ATCC 6538 (11,206 ± 0.342 mm to 750 mg/mL); *A. baumannii* ATCC 19606 (10.570 ± 0.535 mm a 250 mg/mL); *P. mirabilis* (7.636 ± 0.196 mm a 250 mg/mL) e *I. orientalis* ATCC 6258 (7.400 ± 0.190 mm a 250 mg/mL). Secondary metabolites such as tannins, quinones, coumarins, phenolic compounds and flavonoids were determined. *Tagetes erecta* L alcoholic extract has antimicrobial activity against clinically relevant strains and may be considered for further studies against multi-resistant microorganisms in the future.

## Introduction

Antibiotics represent one of the most successful forms of therapy in medicine. But the efficiency of antibiotics is compromised by a growing number of antibiotic-resistant pathogens.^1^ Multidrug resistant patterns in Gram-positive and -negative bacteria have resulted in difficult-to-treat or even untreatable infections.^2^ Widespread resistance to antibiotics among bacteria is the cause of hundreds of thousands of deaths every year.^3^

In February 2017, in light of increasing antibiotic resistance, the WHO published a list of pathogens that includes the pathogens designated by the acronym ESKAPE to which were given the highest “priority status” since they represent the great threat to humans.^4^ The ESKAPE pathogen is a class of drug-resistant pathogens that comprises *Enterococcus faecium* (E), *Staphylococcus aureus* (S), *Klebsiella pneumoniae* (K), *Acinetobacter baumannii* (A), *Pseudomonas aeruginosa* (P), and *Enterobacter* species (E).^5^

The global threat of AMR calls for the collaborative action for developing effective strategies in combating.^6^ Recently, secondary plant metabolites have been used for the management of MDR pathogens.^7^ Numerous bioactive compounds derived from plants, called phytochemicals, have been evaluated, which are comparatively safer than synthetic alternatives and exert multiple therapeutic benefits associated with their high efficacy, for example as antimicrobial.^8^

*Tagetes erecta*, is an herbaceous plant of the family *Asteraceae* native to Mexico and found in other tropical, subtropical areas such as America, India and Bangladesh^9^. It is commonly known as Cempasúchil, is a member and has several names such as Holigold, garden marigold, Marybud, Ganda, among others.^10^ Biochemical studies show that the leaves and flowers of *Tagetes erecta* are rich in alkaloids, flavonoids, tannins and essential oils. These active ingredients have a very important antibacterial activity.^11^

The flowers of *Tagetes erecta* with different solvents show antimicrobial activity against *Escherichia coli, Pseudomonas aeruginosa, Alcaligenes faecalis, Bacillus cereus, Campylobacter coli, Proteus vulgaris, Klebsiella pneumoniae, Streptococcus mutans* and *Streptococcus pyogenes*.^12^

## METHODS

### Plant material and extraction

The botanical specimen was collected in the municipality of Tenango del Valle, State of Mexico. To obtain the extract, a maceration was carried out, where 75 g of the plant (stem, leaves and flower petals) were used in 1250 mL of 70% ethanol and allowed to stand for 14 days. The resulting product was filtered and concentrated at reduced pressure with the Rotavapor (Hahn Shin Scientific, HS-2000NS). Finally, the extract obtained was stored in a cooling system at 4 ºC. The plant was taxonomically identified in the IMSS Medicinal Herbarium, Mexico.

### Phytochemical screening

The identification of secondary metabolites present in the alcoholic extract of *Tagetes erecta* was carried out by colorimetric tests, as follows: saponins (foam), alkaloids (Dragendorff), triterpenes (Rosenthaler), steroids (Rosenthaler), cumarines (Baljet), Quinones (NaOH), Tannins (FeCl_3_), Phenolic compounds (FeCl_3_).^13,14^

### Antibacterial activity by the disc diffusion method^15^

We evaluated 13 microorganisms obtained from the ceparium of the Microbiology laboratory of the University of Ixtlahuaca CUI, which were: 9 bacteria (*Salmonella enterica* ATCC 14028, *Shigella flexneri* ATCC 12022, *Acinetobacter baumannii* ATCC 19606, *Escherichia coli* ATCC 11229, *Pseudomonas aeruginosa* ATCC 27853, *Staphylococcus aureus* ATCC 6538 methicillin-sensitive, *Staphylococcus aureus* ATCC 43300 methicillin-resistant, *Klebsiella pneumoniae* and *Proteus mirabilis*); 4 yeasts (*Candida albicans* ATCC 10231, *Issatchenkia orientalis* ATCC 6258, *Candida glabrata* ATCC 15126, *Candida tropicalis* ATCC 13803).

To confirm the purity of the strains, first they were sown in the general media: Soya Trypticaine Agar (DIBICO) and Dextrose Sabouraud Agar (DIBICO), and then a Gram staining was performed, and selective and differential media were used, In addition to the use of biochemists.

From the extract obtained by alcoholic maceration, different concentrations were prepared with Dimethylsulfoxide (DMSO) (EMSURE ACS), which were: 100 mg/mL, 250 mg/mL, 500 mg/mL, 750 mg/mL and 1000 mg/mL.

Bacterial strains were reembrassed in Soya Tripticaine (TSA) agar (DIBICO®) at 37°C for 24 h and after growth turbidity was adjusted to 0.5 McFarland used sterile isotonic saline solution (0.85% NaCl) to obtain a concentration between 1 to 2 × 10^8^ CFU/mL, where the concentration was corroborated by reading between 0.08 and 0.13 absorbance in the visible range spectrophotometer and UV 30% (Velab®) at a wavelength of 625 nm. After adjusting the strain concentration, sterile swabs were placed in closed straws in Müeller-Hinton agar (DIBICO) boxes under aseptic conditions and 6 mm AA grade sterile discs (Whatman^TM^, Cytiva®) and 10 μL of the concentrations of *Tagetes erecta* alcoholic extract to be evaluated were placed on the discs.

The boxes were incubated at 35 2°C for 20h and halos generated were measured with the help of a vernier. Each test was performed in triplicate. The results of antibacterial activity were analyzed using the statistical program SPSS^16^, by means of an ANOVA test [Analysis of variance; Tukey (p 0.05)]. Discs impregnated with 10 μL of sterile DMSO were used as negative control, which was applied to the culture medium and confirmed that it will not generate halo of inhibition, in addition amikacin 30 μg (BBL^TM^), chloramphenicol 30 μg (BBL^TM^), linezolid 30 μg (BBL^TM^) and Cefoxitine 30 μg (BD BBL^TM^) discs to corroborate methicillin resistance.

Regarding the antimicrobial activity of the yeasts^17^, a suspension was performed with sterile isotonic saline and yeast colonies reembrayed in Dextrose Sabouraud agar for 24 hours; to match McFarland’s 0.5 standard and have a 70% transmittance reading in the spectrophotometer at 530 nm (this solution has an approximate concentration between 1×10^6^ and 5×10^6^ CFU/mL). Subsequently followed the same as in antimicrobial activity bacteria, with the exception that the procedure was performed on Mueller Hinton agar boxes supplemented with 2% glucose and 0.5 μg/ml of methylene blue. As a positive control, discs containing 10 μL of 25 μg/mL Fluconazole solution (Sigma-Aldrich) were used.

## RESULTS

### Identification of plant material

The following table shows the result of the taxonomic identification of the botanical specimen performed by the IMSSM Medicinal Herbarium.

### Phytochemical screening

The secondary metabolites identified in the ethanolic extract of *Tagetes erecta* L are shown in Table 2.

**Table 1.** Taxonomic identification of botanical material.

| Botanical material |  | Institution | Registration number |
| --- | --- | --- | --- |
| Common name | Scientific name |  |  |
| Marigold | *Tagetes erecta* L | Medicinal Herbarium of Mexico IMSSM | 17 098 |

**Table 2.** Identification of secondary metabolites present in the ethanolic extract of *Tagetes erecta* L by colorimetric tests.

| Test | Ethanolic extract |
| --- | --- |
| Saponins (Foam) | - |
| Alkaloids (Dragendorff) | - |
| Triterpenos (Rosenthaler) | - |
| Steroids (Rosenthaler) | - |
| Coumarins (Baljet) | + |
| Quinones (NaOH) | + |
| Tannins (FeCl<sub>3</sub>) | + |
| Phenolic compounds (FeCl<sub>3</sub>) | + |
| Flavonoids (Shinoda) | + |

(+): Presence; (-): Absence (qualitative test).

### Antibacterial activity by the disc diffusion method

Antimicrobial activity with ethanolic extract of *Tagetes erecta* L in the 13 strains analyzed only in 4 strains were observed the presence of inhibition halos, which were: *Acinetobacter baumannii* ATCC 19606 (10.570 ± 0.535 to 250 mg/mL); *Proteus mirabilis* (7.636 ± 0.196 to 250 mg/mL); *Staphylococcus aureus* ATCC 43300 (19.400 ± 0.435 to 500 mg/mL); *Staphylococcus aureus* ATCC 6538 (11.206 ± 0.342 to 750 mg/mL) and *Issatchenkia orientalis* ATCC 6258 (7.400 ± 0.190 to 250 mg/mL). The results of antimicrobial activity in bacteria are shown in Table 3 and Figure 1.

**Table 3.**
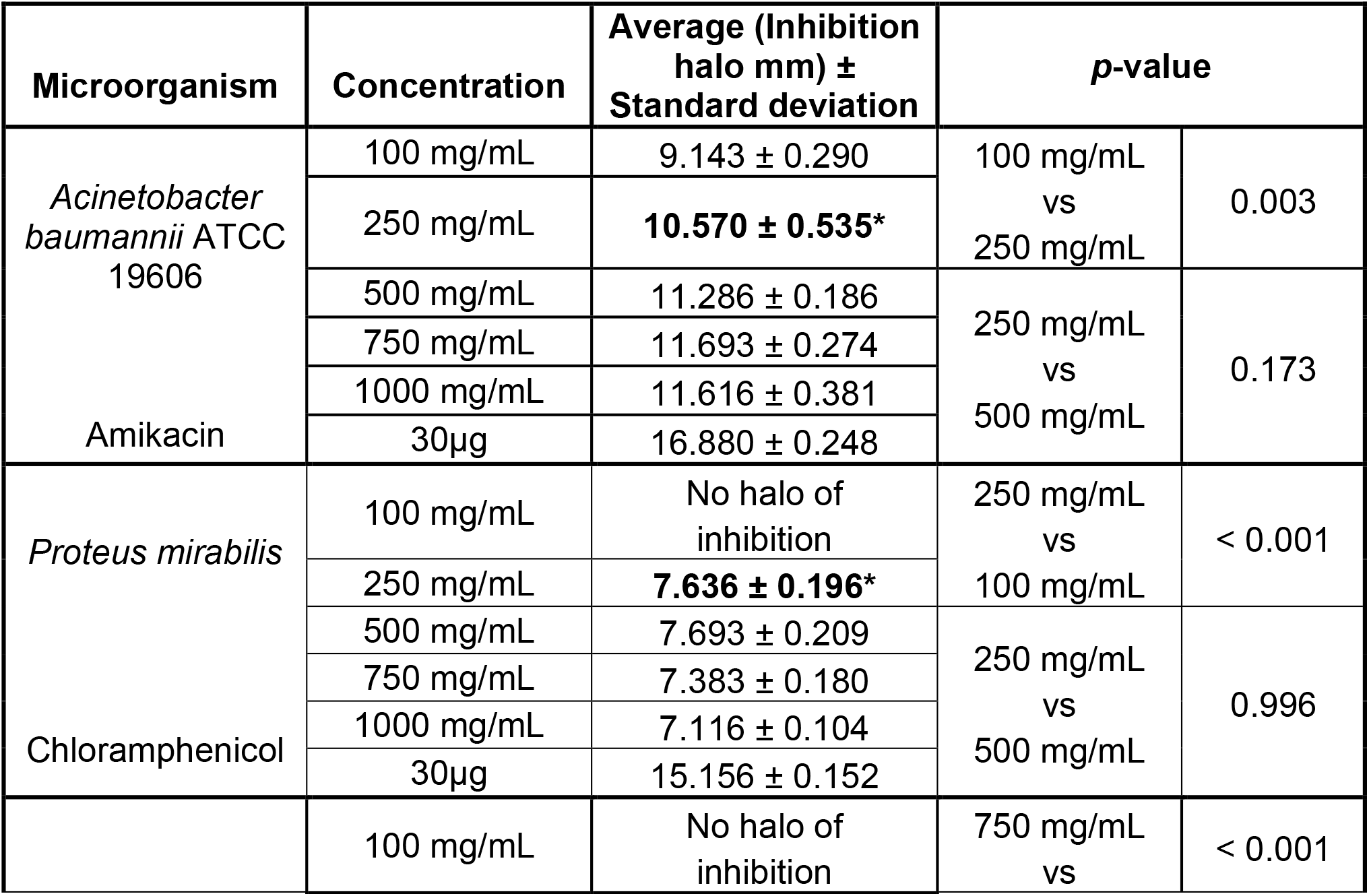
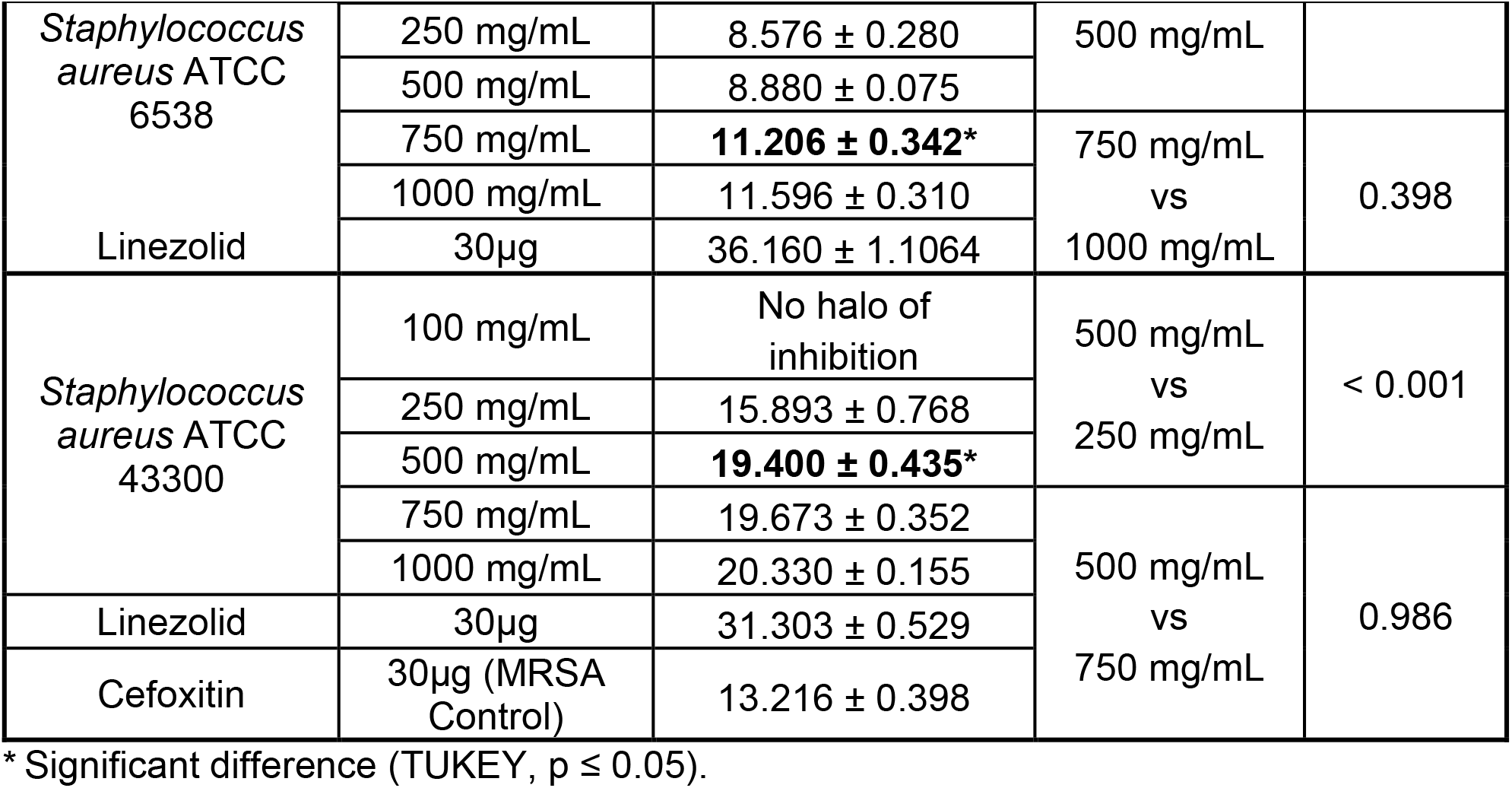
Antibacterial activity of the ethanolic extract of *Tagetes erecta* L against different bacterial strains.

**Figure 1.**
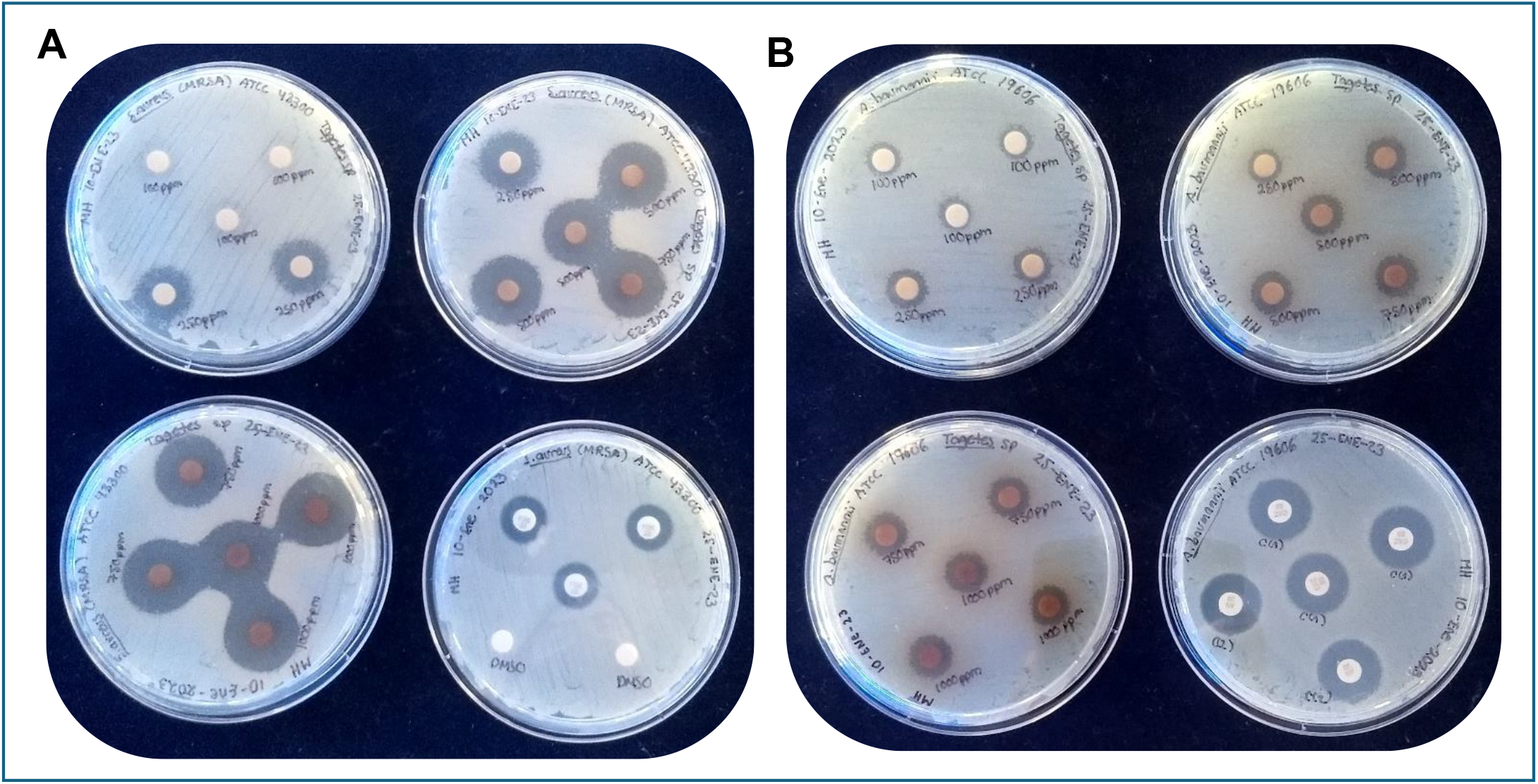
Antimicrobial activity of the ethanolic extract of *Tagetes erecta* L at different concentrations against *Staphylococcus aureus* ATCC 43300 methicillin resistant (left) and *Acinetobacter baumannii* ATCC 19606 (right). Source: Own.

The alcoholic extract of *Tagetes erecta* L showed no antimicrobial activity against the following strains: *Salmonella enterica* ATCC 14028, *Shigella flexneri* ATCC 12022, *Escherichia coli* ATCC 11229, *Pseudomonas aeruginosa* ATCC 27853, *Klebsiella pneumoniae, Candida albicans* ATCC 10231, *Candida glabrata* ATCC 15126 and *Candida tropicalis* ATCC 13803.

## DISCUSSION

The evaluated botanical specimen was identified by the Medicinal Herbarium of Mexico IMSSM-Mexico as *Tagetes erecta* L. In Mexico the plant is known as Cempasúchil, where it is cultivated because of its yellow or orange flowers that are traditionally used during the festivities of the Day of the Dead. There are more than 53 species of this plant worldwide, such as: *Tagetes lucida* Cav., *Tagetes laxa* Cabrera, *Tagetes riojana* M. Ferraro, *Tagetes micrantha* Cav., *Tagetes minuta* L., among others. The plant is used in folk medicine to treat various diseases. Some species of *Tagetes* have been shown to have antimicrobial, anti-inflammatory, hepatoprotective, healing, insecticidal, analgesic properties that are attributed to the presence of secondary metabolites.^9,18^

The presence of tannins, quinones, coumarines, phenolic compounds and flavonoids was determined by phytochemical screening. The results coincide with several works ^18,19, 20^ with the presence of these, although there is the difference that they determined the presence of saponins, alkaloids, triterpenes and steroids, possibly to factors such as: the time of sampling or genotype of samples or method of extraction.

The presence of identified secondary metabolites in the alcoholic extract of *Tagetes erecta* may be contributing to the antimicrobial activity observed as documented^21^, that tannins and flavonoids are capable of forming weak bonds with the membrane proteins of bacteria (such as adhesins) and thereby inactivating their membrane and transport proteins; whereas phenolic compounds and flavonoids when they bind to the bacterial membrane cause alteration and lysis; likewise, tannins and phenolic compounds, alter the metabolism of bacteria involved in cell death by binding to intra- or extracellular protein structures that causes inhibition of oxidative phosphorylation or production of adenosine triphosphate (ATP).

Regarding coumarins, such as scopoletin, umbelliferone, among others, it has been documented^22^ that they are able to inhibit the growth of both bacteria and fungi. In one work, different hydroxycumarins derivatives were synthesized and determined a higher antimicrobial activity in *Staphylococcus aureus* ATCC 6538, establishing that the formation of hydrogen intermolecular bonds can maintain a suitable configuration to bind an enzyme and thus be an important factor in the antimicrobial and antioxidant activities of compounds.^22^

The antimicrobial activity of *Tagetes erecta* extract was shown to be more antimicrobial against methicillin resistant *Staphylococcus aureus*, a clinically relevant strain with a multidrug resistance pattern (penicillin, macrolides, fluoroquinolones, aminoglycosides, tetracyclines and lincosamides). This strain is responsible for the death of almost 50,000 individuals each year in the US and Europe alone. ^23,24^

The inhibition halos obtained for *Staphylococcus* strains were *S. aureus* ATCC 43300 with 19.40 ± 0.43 mm at 500 mg/mL and 11.20 ± 0.34 mm at 750 mg/mL for *Staphylococcus aureus* ATCC 6538. Compared with other works, the following results are obtained: *S. aureus* ATCC 25923 12.3 ± 0.04 mm (leaf) and 19.5 ± 0.14 mm (flower) at a concentration of 200 mg/mL^19^; *S. aureus* ATCC 25923 12 mm (bud) at a concentration of 300 mg/mL^25^; *S. aureus* ATCC 29737 12 mm (flowers) at a concentration of 20 mg/mL^20^. From the reported work mentioned above, the results differ in inhibition halos and concentrations used, possibly due to working with a different solvent (methanol), the part of the plant employed, the method of extraction of metabolites, the difference in composition of secondary metabolites of botanical specimens that differ by region (Cuba, India and Turkey) or by strain used.

*Acinetobacter baumannii* is another important nosocomial microorganism, which has intrinsic and acquired antimicrobial resistance, as well as resisting desiccation, allowing it to survive on inert surfaces for long periods of time and in recent decades has had an increasing resistance to carbapenems, depriving them of these first-line antimicrobial agents to treat A. baumannii infections.^26^ With respect to the result obtained, was observed a halo of inhibition of 10.57 ± 0.53 mm at 250 mg/mL and compared with the reported work done in the School of Biosciences and Technology in India found that of all evaluated strains *A. baumannii* had the greatest antibacterial effect with a activity rate of 0.91333, the work performed.^27^

Of the remaining microorganisms with antimicrobial activity, *P. mirabilis* was observed to have a halo of inhibition of 7.63 ± 0.19 mm at 250 mg/mL, having a value lower than the reported value^20^ at a concentration of 20mg/mL; whereas for the yeast *I. orientalis* ATCC 6258 a halo of 7.40 ± 0.19 mm at 250 mg/mL was determined and reported results^20,25^ mainly report *Candida albicans*, where there is no halo of inhibition and where they report a halo of inhibition between 11 and 15 mm at a concentration of 500 to 1000 μg/mL^28^

Of the thirteen microorganisms evaluated, 2 were Gram bacteria (+) and 7 Gram bacteria (-), and only two of the latter showed a halo of inhibition, thus corroborating several studies^19,21^, which relate the structure of the microorganism that has a layer of polysaccharides in the external membrane of the bacterium that causes a strong protective function and may constitute an obstacle to the entry of water-soluble secondary metabolites into the cell, which are responsible for the antibacterial action.

## CONCLUSIONS

There is a significant increase in the search for natural products with antimicrobial capabilities, which represent a strategy to discover new molecules against multiresistant strains and thus have more therapeutic options. In the work carried out with the ethanolic extract of Tagetes erecta, the presence of tannins, quinones, coumarins, phenolic compounds and flavonoids was determined, and antimicrobial activity was evidenced in three genera of bacteria and one genus of yeast of clinical relevance.

« Conflicts of interest: none»

